# Perceived urban microhabitat heterogeneity impacts carabid beetle communities

**DOI:** 10.1101/2025.10.21.683650

**Authors:** Basile Finand, Meeri Tahvanainen, Heikki Setälä, D. Johan Kotze

## Abstract

**Context:** The habitat heterogeneity hypothesis predicts that increased environmental heterogeneity promotes biodiversity by providing more niches. However, few studies have examined whether human perceptions of heterogeneity align with the ecological responses of arthropods.

**Objectives:** We investigated how perceived habitat heterogeneity translates into environmental variables. Then, we explored the effects of perceived habitat heterogeneity in remnant urban forests and parks on carabid beetle species richness, composition, and functional traits.

**Methods:** We examined carabid beetle (Coleoptera, Carabidae) communities across three habitat types differing in perceived microhabitat heterogeneity: heterogeneous urban forests, homogeneous urban forests, and urban parks in Helsinki, Finland.

**Results:** Environmental data confirmed distinct habitat characteristics: for most variables measured, heterogeneous forest sites were more variable compared to homogeneous forest sites. Contrary to expectations, species richness was higher in homogeneous forests and parks and lower in heterogeneous forests. Community composition differed among habitats and was influenced by canopy openness and dead wood. Heterogeneous forests harboured fewer macropterous and open-habitat species, while homogeneous forests had more brachypterous and generalist species, indicating differences in dispersal capacity and habitat preferences. Body size at the community and population levels, and mass at the population level were generally unaffected.

**Conclusions:** These results suggest that while habitat heterogeneity shapes community structure and traits, species richness does not always increase with heterogeneity, highlighting complex relationships between environmental variables and beetle communities. We underscore the importance of considering specific habitat features, such as dead wood and canopy cover, for urban biodiversity conservation.

## Introduction

Habitat heterogeneity is an important predictor of biodiversity changes following the “habitat heterogeneity hypothesis” (MacArthur & MacArthur 1961; Tews et al. 2004; Stein et al. 2014). Heterogeneity increases resource availability and creates niches, allowing a more diverse set of species to survive (Hutchinson 1957), evidenced in, for instance, such diverse examples as freshwater invertebrates (Taniguchi & Tokeshi 2004), oribatid mites in the soil (Hansen 2000), and tropical mammal communities (August 1983). However, this pattern is not universal (see Bartholomew et al. 2016). Heterogeneity can be created by various anthropogenic factors, such as habitat fragmentation, which is increasing edge effects, microclimatic variation, or patch connectivity (Fahrig 2003). In an urban forest context, management practices have an important influence on habitat heterogeneity, especially for arthropods (Benton et al. 2002). For instance, logging and management will alter forest structure, canopy cover, and deadwood availability (Hall et al. 2003; Cazzolla Gatti et al. 2015). The presence of this deadwood, for instance, will determine the availability of habitat for many species (Lassauce et al. 2011; Siitonen 2001; Seibold & Thorn 2018). Deadwood will also alter microclimatic conditions in its vicinity, for instance an increase in humidity and lowered temperatures. Additionally, the use of urban forest by humans creates areas of compact soils due to walking paths and increases soil pollution. Together, these human actions create a plethora of soil micro-conditions, enhancing habitat heterogeneity. Such heterogeneity can, however, create a trade-off between an increase in species richness due to habitat heterogeneity and its decrease due to the loss of suitable habitat area (Allouche et al. 2012). Our question, therefore, is whether this heterogeneity in urban greenspace translates into a more diverse and distinct arthropod community as compared to less heterogenous urban greenspace, which, if so, will have management and planning implications.

The carabid beetle taxon (Coleoptera, Carabidae) is a well-studied group of arthropods. They are commonly used in ecological studies due to their sensitivity to environmental change and habitat alteration (Lövei & Sunderland 1996; Koivula 2011; Kotze et al. 2011; Rainio & Niemelä 2003). These beetles are also diverse in terms of their traits, including body size, trophic level, degree of habitat generalism, feeding strategy and dispersal capacities (Lindroth 1985, 1986), allowing a nuanced understanding of the response of communities to environmental and habitat change. Several studies have demonstrated the importance of micro-environment characteristics on assembling carabid beetle communities. For instance, litter depth, soil moisture content, tree and shrub density, the presence of logs, canopy openness, or soil pH have been shown to affect carabid beetle diversity and assemblages (Paje and Mossakowski 1984; Koivula. et al. 1999; Janssen et al. 2009; Barton et al. 2009; Ogai and Kenta 2016; Thorn et al. 2016). Additionally, community parameters including abundance, richness and/or diversity have been shown to be higher in structurally complex forests (Lassau et al. 2005), in heterogeneous habitats (Marrec et al. 2021), and the heterogeneity along an urbanisation gradient (Magura et al. 2004). Therefore, local variation of these characteristics, which creates habitat heterogeneity, is key in explaining local carabid beetle diversity. Communities can vary not only in their taxonomic diversity but also in their functional diversity. Heterogeneity affects the functional diversity of carabid beetles with, for instance, earlier breeding species dominant in simple open habitat (Duflot et al. 2014). Moreover, larger and apterous (without wings) species are more commonly found in woody heterogeneous landscapes, while small and macropterous species are found in open landscapes (Kędzior & Kosewska 2022).

Habitat heterogeneity can be assessed visually by estimating the quantity of deadwood, different layers of vegetation, or canopy cover differences, with a number of studies investigating the response of carabid beetles to habitat heterogeneity (Blanchet et al. 2013; Staudacher et al. 2018). However, it is not clear how this human-perceived heterogeneity is related to actual measurable environmental characteristics and how it impacts biodiversity, especially in the urban milieu. The aim of this study was to investigate how perceived habitat heterogeneity, used by the authors as a proxy of structural complexity of the habitat, translates into measurable environmental variables. Then, we explored the effects of this perceived ground-level habitat heterogeneity and the related environmental variables in remnant urban forests and parks on carabid beetle species richness, composition, and functional traits, such as body length and type of wings. Finnish cities provide a great opportunity to investigate this, since they consist of remnant forests that are differently used and managed, thus creating homogeneous and heterogeneous undergrowth and varying levels of complexity of woody debris. For instance, dead wood is kept in some forests, while it is removed from others. The dominant tree species, Norway spruce (*Picea abies*), is common to most urban forests irrespective of their disturbance level and can also be found in urban parks with a highly homogenous undergrowth. We hypothesise that habitat heterogeneity has a significant effect on carabid beetle species composition, with homogenous (managed) sites consisting of more generalist and open-habitat species, while heterogeneous (more natural) sites are characterised by a larger proportion of forest species (McKinney 2006; Gaublomme et al. 2008). We predict a higher beetle diversity in heterogeneous sites compared to homogeneous ones due to the higher number of different niches. However, since open-habitat genera in the carabid beetle family are species-rich (Lindroth 1985, 1986, see also references in Venn & Kotze 2014), this could counterbalance the effect of heterogeneity with more species in more open homogeneous sites. Additionally, these communities are expected to consist of a higher proportion of small and flight-capable species compared to those in heterogeneous sites. We also hypothesise larger and brachypterous species in the forest sites (Kędzior and Kosewska 2022). Last, we looked at the impact of heterogeneity on intraspecific traits such as body size and mass. If structural homogenisation at ground level places these beetles under higher stress in terms of reduced resources and increased predation pressure, we hypothesise that beetles in homogenous sites will be smaller in body size and lighter in mass.

## Material and methods

### Study area and sites

The study was conducted in the City of Lahti, southern Finland (60°58′55″ N, 25°39′32″ E, Figure 1a). Altogether, 21 sampling sites were selected within the city, which is located in the southern boreal vegetation zone (Table S1). The sites were divided into three different habitat types (7 replicates per type): perceived heterogeneous (HET, Fig. 1b) and homogeneous (HOM, Fig. 1c) spruce-dominated, *Myrtillus* type urban remnant forests, and below an individual spruce tree in highly homogeneous urban parks (PARK, Fig. 1d). The forests investigated are more than 80 years old and experience minimal management. The size of the heterogeneous forests ranged between 7.1 ha and 16.9 ha, and between 7.5 ha and 16.9 ha for the homogeneous forests (one homogeneous HOM4 and one heterogeneous HET4 site, not included here, were situated in a huge continuous forest of more than 100 ha). Parks were smaller and ranged between 0.04 and 11.0 ha. Selecting the forest sites was implemented by visually evaluating the habitat. The main indicator for heterogeneity in this study was the amount of dead and decaying wood, while homogenous sites had very little or no dead wood.

**Fig. 1:**
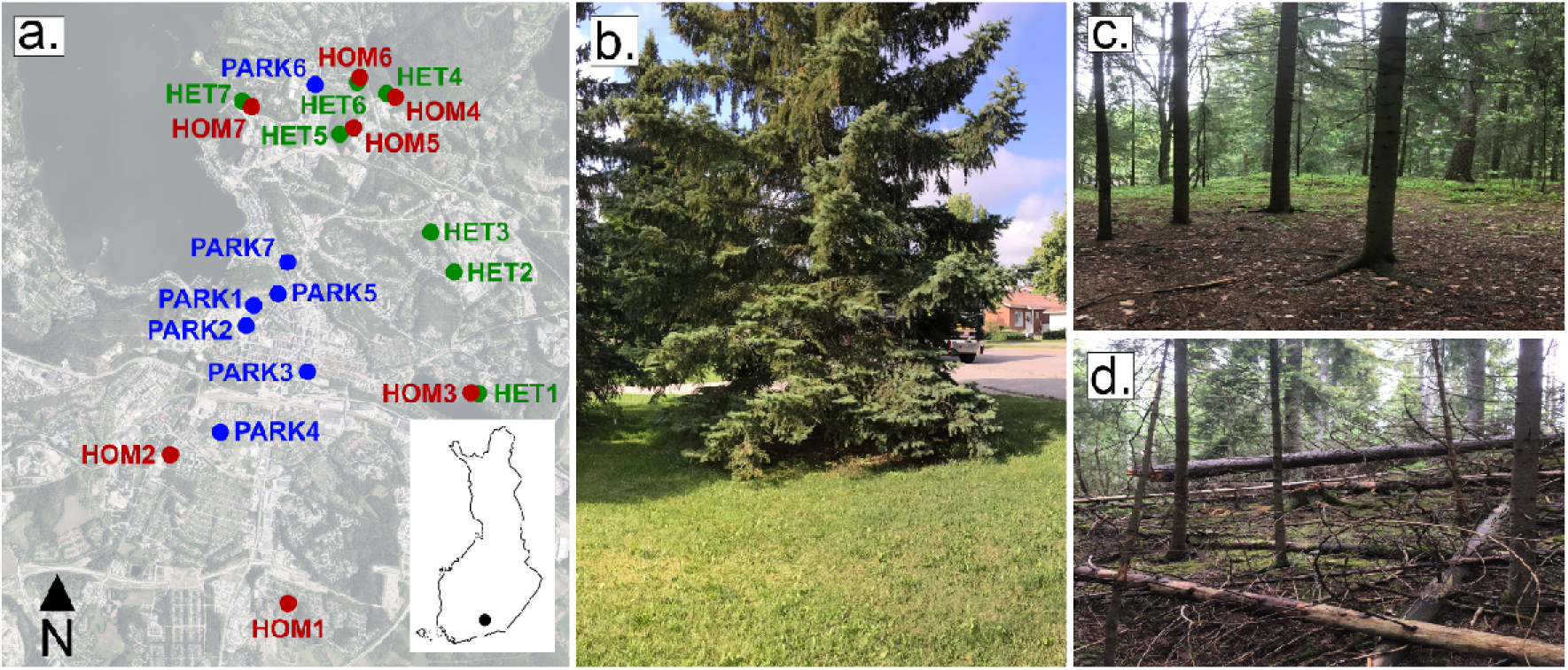
a) Map of the sites with the three site categories: parks (blue), homogeneous forests (red), heterogeneous forests (green). The insert map in the bottom right corner indicates the location of the city of Lahti in Finland. b) c) and d) photos of respectively a park (PARK), a homogenous forest (HOM) and a heterogeneous forest (HET).

### Carabid beetle sampling

Carabid beetles were sampled by continuous pitfall trapping in 2021 from the 31^st^ of May to late 21^st^ of October. Placing pitfall traps in the environment is a passive way to collect specimens and is affected by the activity of the beetles in their habitat, as well as the size of the population (Lövei and Sunderland 1996). The traps were plastic cups (depth and mouth diameter 6.5 cm) that were dug into the ground, their rim flush with the ground surface. Each cup was half-filled with a 50 % propylene glycol–aqueous solution to preserve the beetles. Above each trap, a 10 x 10 cm plastic roof was placed at about 2 cm above ground to protect the traps from rain, excessive debris and possible disturbance caused by small mammals ending up in the traps or eating the catch.

In the park sites, pitfall traps were placed directly under a conifer canopy to avoid possible disturbance caused by passers-by. Unlike park traps, forest traps were placed somewhere in the middle of a forest patch that filled the criteria for a heterogeneous or homogeneous remnant forest. Five traps were installed at each site (105 traps altogether) and placed within a square meter, with four traps placed at the square’s corners and the fifth trap in the middle of the square. The catch of the five traps per site was pooled per visit.

Samples were collected every third week, resulting in seven visits. The catch of all five traps per site and across all visits was pooled for the analysis. All trap losses, along with other noteworthy remarks about the condition of each site, were recorded and considered in data analysis to standardise the number of beetles by trapping day. The collected catches were preserved in denatured alcohol before sorting and identification. From each sample, the number of carabid beetles was recorded. Each individual was identified to species level with the identification keys in Lindroth (1985, 1986) and supporting online materials such as *laji.fi* and *kerbtier.de*. Identification was conducted using a Leica S4E stereo-zoom microscope (Table S2).

Beetle traits were measured from each individual collected. Body length, elytra length and dry body mass (air-dried in the laboratory for seven days before weighing on an analytical balance; precision at 0.0001 g) were measured for each carabid beetle collected. Elytra length was chosen to represent body length since it is easier to measure accurately than the full body length of a beetle, with body length and elytra length being highly correlated (Kotze et al. 2024). We corrected the mass per size to have the relative change of mass independently of the change of size (calculated as mass/size). Wing type (brachypterous = reduced wings, macropterous = full wings) was also recorded from the collected individuals. These direct measurements were also used in the species-level analyses. Traits at the species level were collected from the literature (Lindroth 1985, 1986), including habitat preference (forest or open habitat specialist, or generalist), food preference (carnivorous, phytophagous or omnivorous), and moisture preference (xerophilic, mesophilic, or hydrophilic).

### Environmental variables

To determine if the sites selected were indeed different in terms of the visual homogeneity/heterogeneity classification in the field, several environmental variables were recorded at each of the 21 sampling sites in mid-August 2021 (Table S1). From a 10 x 10 m square around the traps, the number of canopy and sub-canopy trees was recorded, along with the volume of dead and decaying wood, such as logs and stumps. From these squares, the area of trampling (all the paths observed as a % of the area of the square) and the presence of anthropogenic items were visually estimated.

In addition to the 10 x 10 m areas, five smaller squares (1 m^2^ each) were placed inside the bigger square – one around the set of traps and four randomly. From these five squares, the percentage vegetation covers (field layer: low-growing vegetation including herbs and grasses, ground layer: vegetation that covers the ground, typically the moss layer, bare ground: no live vegetation), the percentage of deadwood, and the closeness of the canopy were visually estimated. Soil litter layer depth (cm) was measured with a nail from three different spots within each of the small squares. To measure soil pH, soil moisture (%) and soil organic matter (%), soil samples were collected from each of the five squares of each sampling site from the top 5 – 10 cm of soil, excluding the litter layer.

Soil samples were stored in a cold room (∼ 5 °C) and analysed during the autumn of 2021. Before analysis, the samples were homogenised with a sieve and a bucket. To avoid contamination, the sieve and the bucket were rinsed and dried between each sample using warm tap water and clean paper towels. Soil pH was measured with an inoLab pH 720 meter from a well-mixed and stabilised soil and distilled water suspension (1:4 (v/v) ratio). Soil moisture content was determined by weighing the soil before and after ∼ 24 h in an oven at 105 °C. These dry samples were placed into a muffle oven (5 h at 550 °C), and the loss on ignition indicated the amount of organic matter. The percentage of moisture and organic matter was calculated by using the before and after oven soil masses. For data analysis, the average of each environmental variable measured from the five squares was calculated to get one value per variable per sampling site.

### Data analysis

Data analyses were performed using R version 4.2.2 (R Core Team 2022).

In order to check that our visual selection separated sites into 3 categories (heterogeneous, homogeneous, and parks) depending on the environmental variables measured, we performed a Principal Components Analysis (PCA). We verified that heterogeneous and homogeneous forests differ in terms of environmental heterogeneity by examining the standard deviation (SD) of variables measured across the five small quadrats within each site, including vegetation cover, deadwood, canopy, litter depth, pH, and organic matter. A high SD indicates substantial variation in a variable among quadrats within the same site, reflecting greater habitat heterogeneity. We performed a linear model per variable to test if the differences are significant. A variable was log-transformed if the residuals of the model were not normally distributed.

To reduce the number of explanatory variables and since many were correlated, we selected the uncorrelated variables by comparing them two by two. When both were numeric, we used a Spearman test. When we had one numeric variable and one factor, we used a Kruskal-Wallis test, while a χ^2^ test was used when both variables were factors. We selected two datasets of variables because the treatment (a three-factor variable including homogeneous and heterogeneous forests, and homogeneous parks) was correlated with many environmental variables. One dataset of variables included only the treatment, and the second dataset of variables included only the uncorrelated variables (without treatment) to evaluate which specific environmental variables were important to the beetle communities. We log-transformed some of the variables to improve their normality: deadwood volume, % trampling, and % bare ground.

In order to observe the impact of treatment and the environmental variables on species diversity, we conducted two analyses. First, we compared species richness between our treatments using a GLM with a Poisson error. We did two separate tests with the two datasets of variables described below. Second, we calculated the cumulative number of species for each treatment using the function *iNEXT* (Hsieh et al. 2024).

To compare community composition between our treatments and the effect of environmental variables, we performed a db-RDA (Distance-based Redundancy Analysis) using the *capscale* function in the *vegan* package (Oksanen et al. 2022). We did two analyses using each dataset of variables. Pairwise comparisons between the three treatments were performed using the *multiconstrained* function in the package *BiodiversityR* (Kindt and Coe 2005).

We used the same procedure for the GLMs and db-RDAs. First, we did the test with all the variables. We checked conditions like normality of the residuals, and overdispersion for the GLMs using the *Shapiro.test* function and the *testDispersion* function of the package DHARMa (Hartig 2022). We calculated the AIC of the model. Then, we removed the non-significant variables one by one until we had the lower AIC. Using this final, most parsimonious model, we observed the statistical significance of the variables.

Additionally, we performed a fourth-corner analysis to assess the relation between beetle traits and the environment (Dray et al. 2014). First, we checked if some traits are correlated using the same tests as for the environmental variable correlations. As some traits were correlated, we decided to keep body size, wing morphology, feeding preference, and habitat preference to have a maximum number of independent variables of ecological significance. Second, we used the *fourthcorner* function of the *ade4* package (Thioulouse et al. 2018) with 10000 repetitions to assess the correlations between traits and environmental variables of the two datasets.

At the population level, we tested for the effects of treatment on elytra size and mass corrected for size of the most represented species. These included species with at least ten individuals collected from at least two treatment categories: *Pterostichus melanarius, P. niger, P. oblongopunctatus, Carabus nemoralis, Patrobus atrorufus, Calathus micropterus, Amara brunnea,* and *Leistus ferrugineus.* If for some species, fewer than ten individuals were measured in one treatment category, we removed this category from the analysis. Because size measurements were performed to the nearest 0.5 mm, data are not continuous and thus not normally distributed, so the non-parametric Kruskal-Wallis rank sum test was used.

For all analyses, a p-value < 0.05 was interpreted as a significant effect, and a p-value < 0.1 as an indicative or slight effect.

## Results

We collected 1637 carabid beetle individuals from 35 species. The most common species were *Pterostichus melanarius* (331 individuals), *Carabus nemoralis* (271), *Patrobus atrorufus* (232), *Calathus micropterus* (206), and *Pterostichus oblongopunctatus* (154) (see Table S2).

The PCA assessing our visual selection of the sites confirmed the separation of sites between the three heterogeneity categories (Fig. 2). The first axis (41.4% of the variation) separated urban remnant forests from the parks. Soil characteristics like a higher pH and field layer percentage, low organic matter content, moisture, and ground layer percentage characterised the parks. The second PCA axis (14.9% of the variation) separated heterogeneous from homogeneous urban remnant forests. A more open canopy, more dead wood, and less trampling were the main factors associated with the heterogeneous sites. Our calculation of variation (standard deviation) showed that most of the variables have greater variability in heterogeneous forests compared to homogeneous ones (Fig. 2b-e). The field layer (ANOVA LM, F = 6.909, p = 0.006), deadwood (F = 10.74, p = 0.007), and organic matter (F = 12.276, p = 0.004) displayed significantly greater variability in heterogeneous forests. Canopy (F = 3.625, p = 0.081), and pH (F = 4.171, p = 0.064) had an indicatively higher variability in heterogeneous forests. The bare (F = 0.22, p = 0.647), and ground layers (F = 0.017, p = 0.898) showed no differences. Finally, litter depth (F = 3.686, p = 0.079) had an indicatively lower variability in heterogeneous forests.

**Fig. 2:**
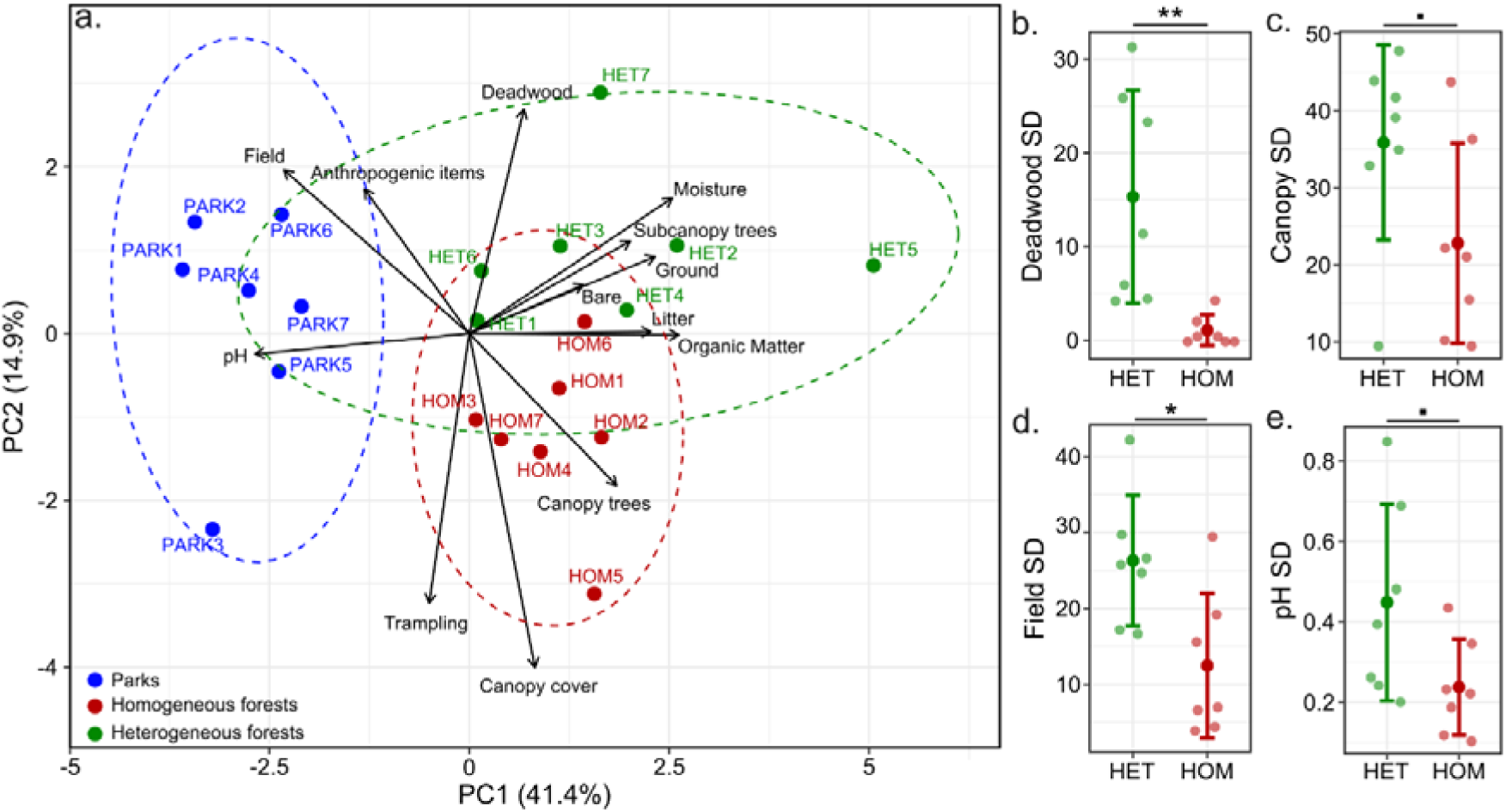
a) Principal component analysis (PCA) of the environmental variables measured in the sites. The percentage of variance is displayed on the x and y axes. Ellipses represent 95% confidence intervals. The mean value of each variable for each site is presented in Table S1. b), c), d), and e) show the standard deviation (SD) of each variable, calculated across the five quadrats within each site, showing higher variability in the heterogeneous forests compared to the homogeneous ones, as expected.

We used two datasets of variables in all of our subsequent analyses. Treatment was correlated with most of the environmental variables measured (Fig. S1). This finding confirmed that our treatments are well separated by the environmental variables measured. Treatment on its own represented the first dataset of variables. To assess which specific environmental variables are correlated with beetle communities, we replaced treatment with trampling area, number of anthropogenic items, canopy cover, bare ground layer, and deadwood, which were all uncorrelated and represent our second dataset of variables.

### Species diversity

Significantly more species were collected from the homogeneous forests compared to the two other treatments (ANOVA GLM, χ^2^ = 9.157, p = 0.01, pseudo-R^2^ = 0.081; pairwise comparison: HOM-HET, p = 0.057; PARK-HET, p = 0.877; PARK-HOM p = 0.016, Fig. 3a). The mean number of species in the heterogeneous forests was 7.86 (±3.58 SD), 11.71 (± 1.60 SD) in the homogeneous forests, and 7.14 (±3.39 SD) in the parks. Parks were characterised by open habitat and generalist species, while both heterogeneous and homogeneous forests were primarily characterised by forest habitat specialists and generalist species (Fig. 3c, d, e). The cumulative number of species was higher in parks (observed = 26, estimated = 38.05) than in homogeneous forests (observed = 22, estimated = 24.66) and heterogeneous forests (observed = 16, estimated = 16) (Fig. 3b).

**Fig. 3.**
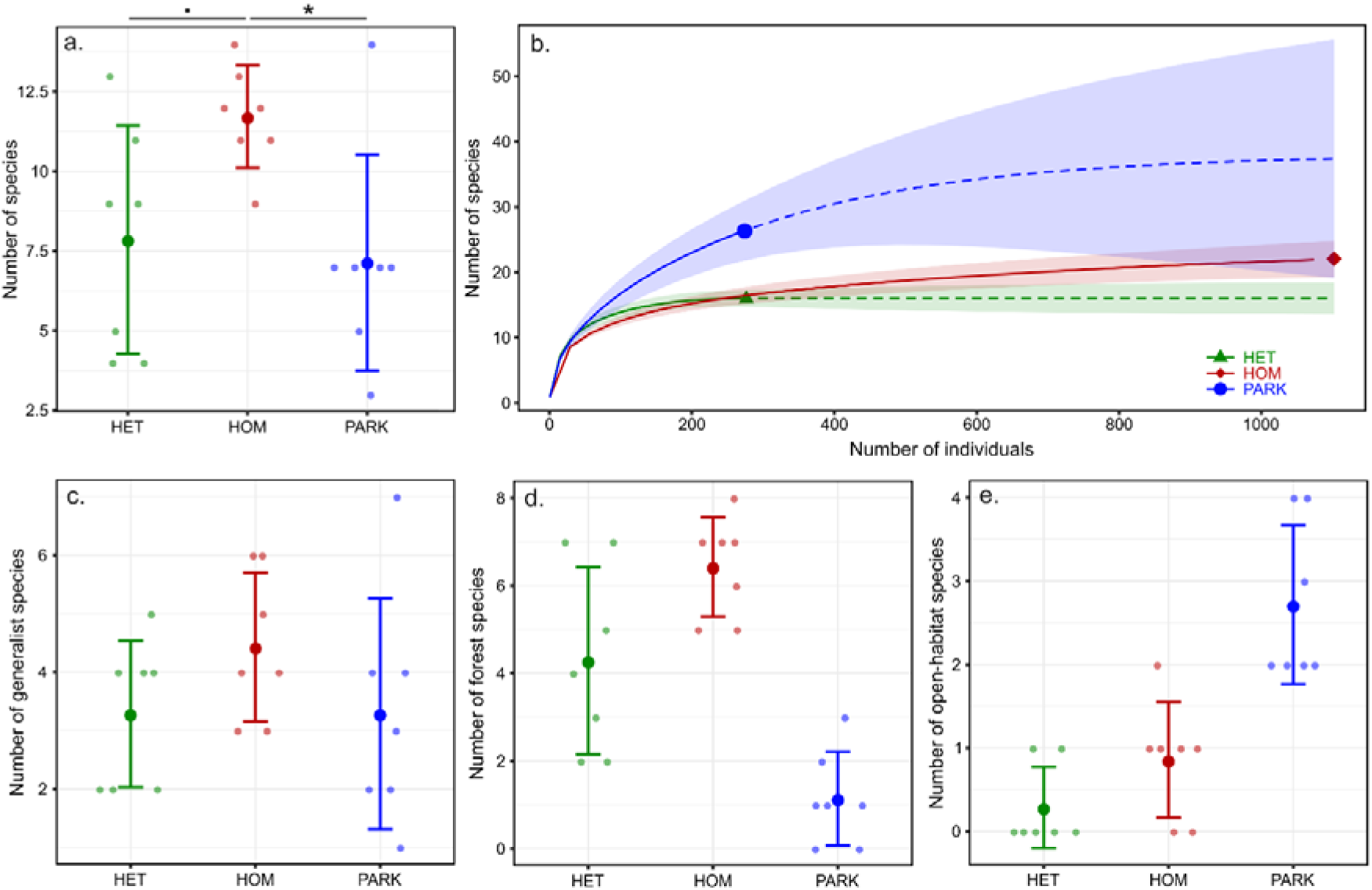
a) Total number of species (± SD) per treatment. The star indicates a significant effect of treatment on the number of species with a p-value < 0.05, and a dot indicates a p-value < 0.1. b) Cumulative number of species per treatment. The plain lines represent the observed number of species. The dashed lines are the estimation of species number for the same sampling effort. c), d), and e) Respectively, the number of generalist species, forest habitat specialists, and open habitat specialists per treatment.

Using the second dataset of variables, carabid beetle richness responded significantly negatively to the number of anthropogenic items (ANOVA GLM, χ^2^ = 4.205, p = 0.04), while slightly positively to trampling and slightly negatively to percentage bare ground (trampling: ANOVA GLM, χ^2^ = 3.148, p = 0.076; bare layer: ANOVA GLM, χ^2^ = 3.322, p = 0.068; pseudo-R^2^ of the model = 0.08).

Carabid beetle community composition was clearly different between treatments (db-RDA, F = 4.068, p = 0.001, constrained explained variation = 0.311; pairwise comparison, HOM-HET p = 0.001, PARK-HET p = 0.002, PARK-HOM p = 0.001) (Fig. 4a). The first axis (20% of the variation) separated park beetle communities from those of remnant forests, and the second axis (11% of the variation) separated communities of heterogeneous forests from the other treatments. Additionally, greater variation in community composition was displayed in the heterogeneous forests compared to the parks, and especially to the homogeneous forests (Fig. 4a). Using the second dataset of variables, deadwood quantity (db-RDA, F = 2.974, p = 0.003) and canopy cover (db-RDA, F = 2.331, p = 0.006) were significant variables in structuring the beetle communities (Fig. 4b). The model explained 34% of the variation. Even though treatment was not in the model, deadwood and canopy cover clearly separated the beetle communities in the parks, the heterogeneous, and the homogeneous forests.

**Fig. 4:**
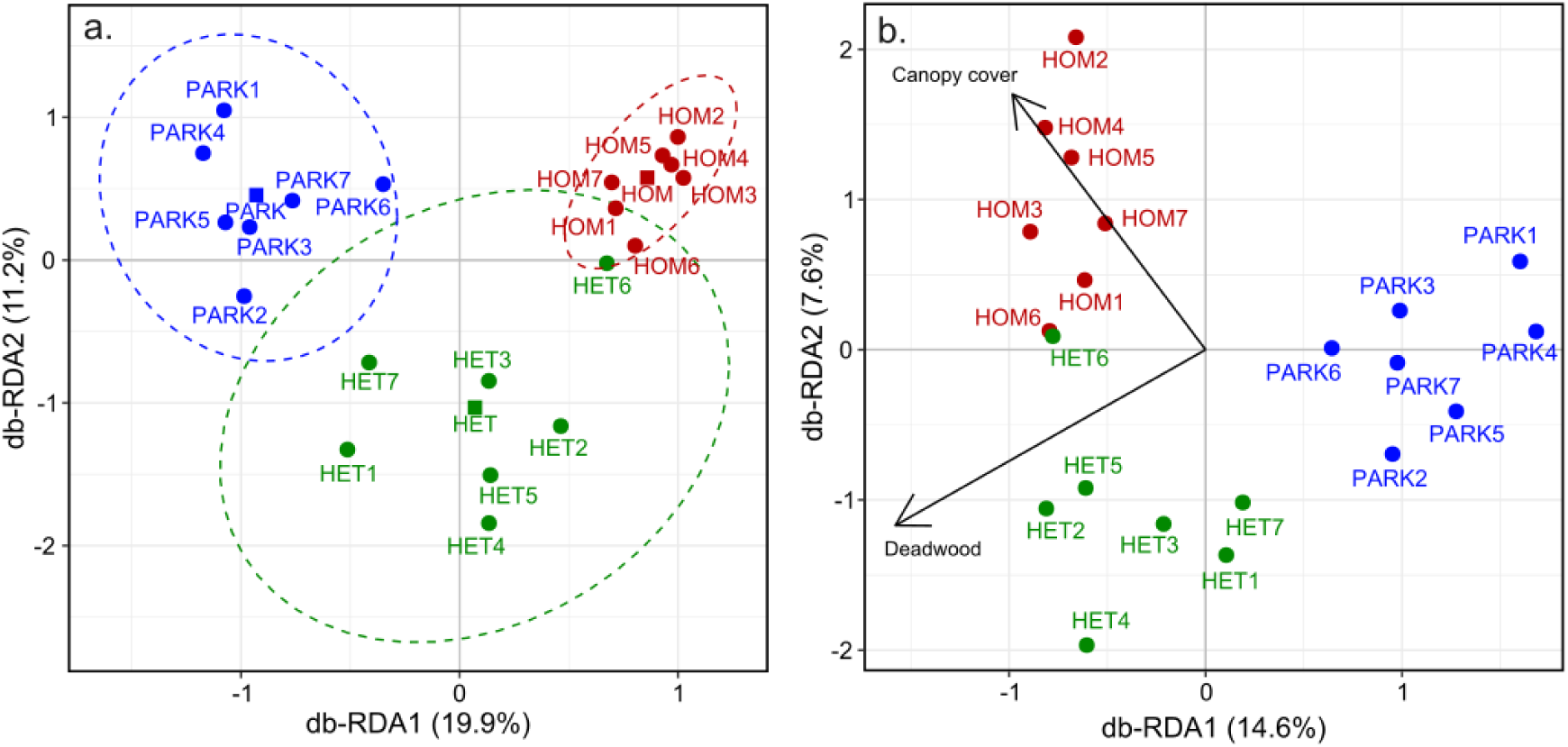
Distance-based Redundancy Analysis (db-RDA) plots with a) the treatment, b) the environmental variables. Percentage variances are displayed on the x and y axes. Ellipses represent 95% confidence intervals.

### Trait analysis at the community level

The trait analysis indicated that heterogeneous forests are characterised by fewer macropterous (p = 0.013) and open-habitat species (p = 0.048). Homogeneous forests also had fewer macropterous species (p = 0.007) and more brachypterous (p = 0.002) and generalist species (p = 0.028). Parks hosted fewer forest-habitat species (p = 0.035) (Fig. 5). When we focused on the environmental variables, sites with more deadwood had more forest-habitat (p = 0.008) and showed an indicative trend towards fewer open-habitat species (p = 0.067). Indicatively, trampling selected open-habitat species (p = 0.09) but fewer generalist species (p = 0.095). The presence of anthropogenic items slightly correlated with more open-habitat species (p = 0.064). Finally, an increase in the proportion of bare ground had an indicatively negative effect on the proportion of generalist species (p = 0.081) (Fig. 5).

**Fig. 5:**
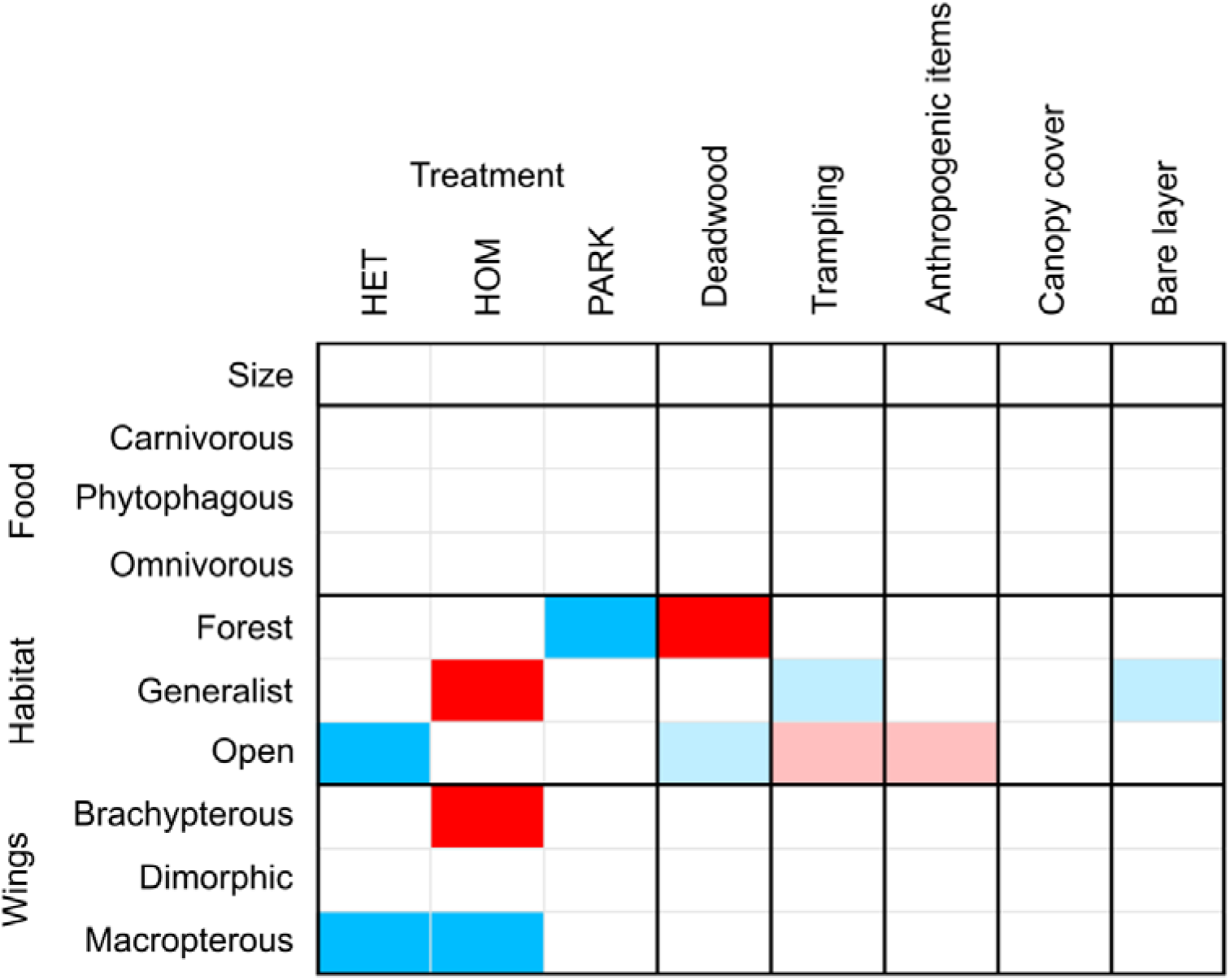
Fourth-corner analysis between species traits and the environmental variables. Dark blue and dark red represent, respectively, a negative and a positive correlation between the trait and the environmental variable with a p-value < 0.05. Light blue and red represent a p-value < 0.1.

### Elytra size and mass at the population level

Eight species had at least ten individuals collected from two treatment categories: *Amara brunnea, Calathus micropterus, Carabus nemoralis, Leistus ferrugineus, Patrobus atrorufus, Pterostichus melenarius, P. niger,* and *P. oblongopunctatus*.

Treatment had no significant effect on the sizes of most of the species evaluated: *P. melanarius, P. niger, Amara brunnea, P. oblongopunctatus, Calathus micropterus,* and *Leistus ferrugineus.* Only *C. nemoralis* (Kruskal-Wallis, χ^2^ = 9.386, p = 0.009; pairwise comparison: HOM-HET, p = 0.01; PARK-HET, p = 0.008; PARK-HOM, p = 0.01), and *Patrobus atrorufus* (Kruskal-Wallis, χ^2^ = 11.665, p < 0.001) were affected by treatment, with *C. nemoralis* generally being smaller in parks and *P. atrorufus* generally being smaller in heterogeneous forests (Fig. 6a).

**Fig. 6:**
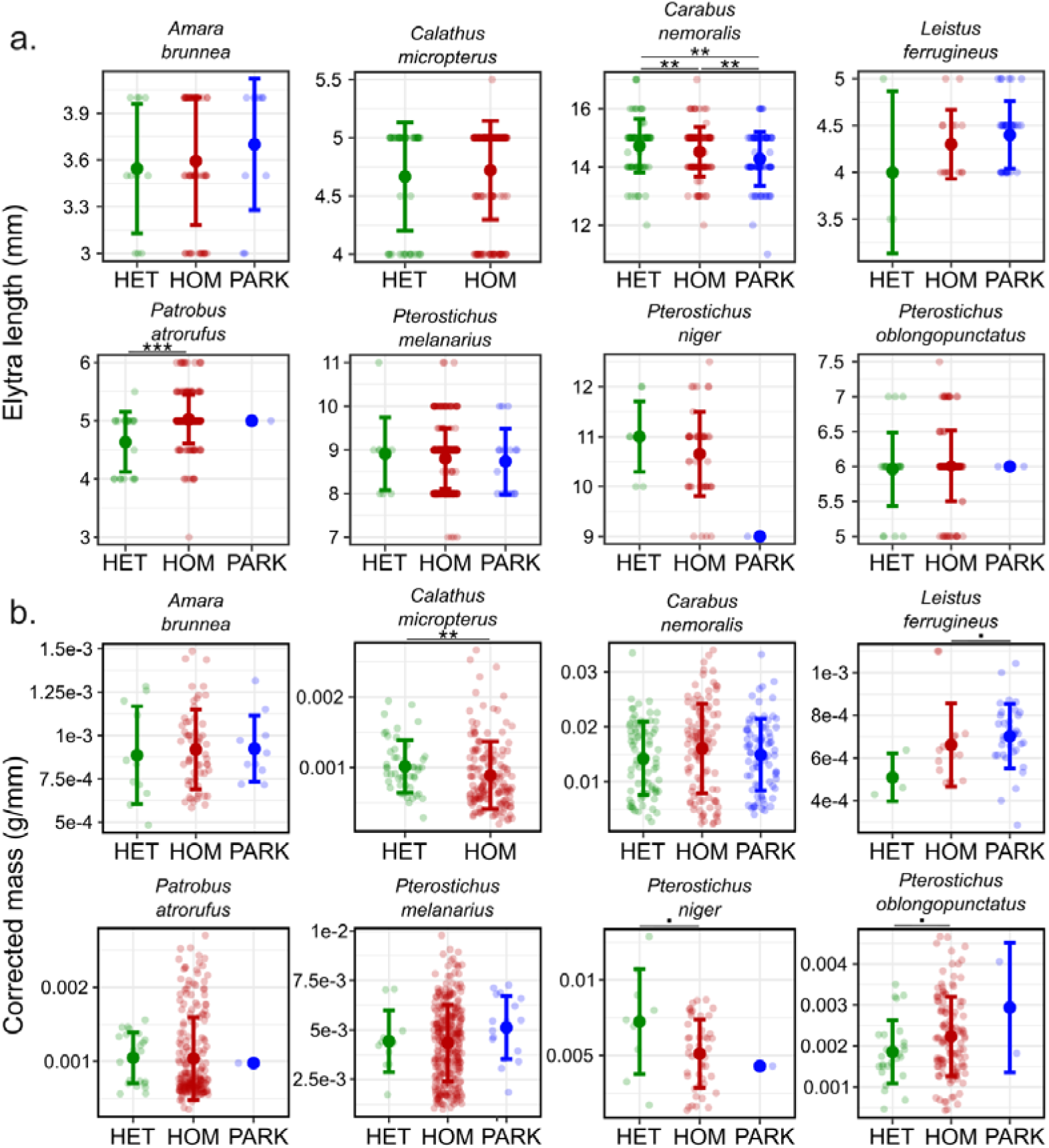
a) Elytra size and b) dry body mass corrected by size across the three treatments. Raw datapoints are shown, as well as mean and SD bars. Three stars indicate a p-value < 0.001, two stars a p-value < 0.01, one star a p-value < 0.05, and a dot a p-value < 0.1. A statistical test has been done only between treatments with more than five individuals.

Concerning beetle mass corrected by size, treatment did not affect *P. melanarius, C. nemoralis, Patrobus atrorufus,* and *Amara brunnea. Calathus micropterus* was heavier in heterogeneous compared to homogeneous forests (Kruskal-Wallis, χ^2^ = 6.699, p = 0.01), *Pterostichus oblongopunctatus* was slightly heavier in homogeneous compared to heterogeneous forests (Kruskal-Wallis, χ^2^ = 3.069, p = 0.08), *Pterostichus niger* was slightly lighter in homogeneous compared to heterogeneous forests (Kruskal-Wallis, χ^2^ = 3.313, p = 0.069), and *Leistus ferrugineus* was slightly lighter in homogeneous forests compared to parks (Kruskal-Wallis, χ^2^ = 3.016, p = 0.082) (Fig. 6b).

## Discussion

We investigated the effect of perceived microhabitat heterogeneity on carabid beetle communities by comparing three urban habitat types: heterogeneous forests, homogeneous forests, and parks. Environmental measurements supported our category selection: parks had higher soil pH and field layer cover, but lower organic matter content, moisture, and ground layer cover. Heterogeneous forests were characterised by a more open canopy, a greater amount of dead wood, and reduced trampling; factors that contribute to increased structural heterogeneity. Contrary to our expectations, species richness was lower in heterogeneous forests than in parks and homogeneous forests due to the influx of open-habitat species into these habitats. Carabid community composition differed markedly among the three habitat types and was mainly influenced by canopy openness and dead wood volume. Parks, despite their high homogeneity, harboured more macropterous species, while homogeneous forests had a higher proportion of brachypterous species compared to heterogeneous forests. At the species level, we found no consistent effect of habitat heterogeneity on individual size or mass. Overall, our findings highlight the role of microhabitat heterogeneity in shaping carabid beetle community composition.

As expected, the categories of perceived heterogeneity were supported by the environmental variables we measured. This confirms that our perception of forest heterogeneity corresponds to actual environmental differences. Our direct quantification of variability, using SD analysis for each variable within each site, showed higher variability in heterogeneous forests for most of the variables, as predicted, indicating greater within-site heterogeneity. This pattern was especially pronounced for key variables of interest, such as deadwood and canopy, but was also evident for soil properties (e.g., pH) and vegetation structure (e.g., field layer). Only one variable exhibited the opposite trend (litter depth), but the difference was marginal. Moreover, additional differences in the direct environmental measurements were observed among our site categories.

Surprisingly, we found a higher average species richness in homogeneous forests compared to both heterogeneous forests and parks. In terms of cumulative species richness, parks hosted the most species overall, followed by homogeneous forests and then heterogeneous forests. These findings do not align with the “habitat heterogeneity hypothesis” (MacArthur and MacArthur 1961; Tews et al. 2004; Stein et al. 2014), which predicts that increased environmental heterogeneity supports a greater number of species by creating more ecological niches. One possible explanation lies in the regional species pool. In Finland, open-habitat species are more numerous than forest specialists (Lindroth 1985, 1986). As a result, parks, although highly homogeneous, may attract a wide range of generalist and open-habitat species, inflating species richness. However, the higher richness observed in homogeneous forests compared to heterogeneous ones is more surprising and contradicts several previous studies on carabid beetles (Lassau et al. 2005; Janssen et al. 2009; Marrec et al. 2021), as well as other taxa such as benthic invertebrates (Taniguchi and Tokeshi 2004), mammals (August 1983), and mites (Hansen 2000). Nonetheless, a similar pattern was observed for darkling beetles (Coleoptera: Tenebrionidae) by Bartholomew et al. (2016). One potential explanation is that canopy closure, which is more pronounced in our homogeneous forest sites, may itself promote higher carabid richness. Thorn et al. (2016) found a positive relationship between closed canopies and beetle diversity. Another possibility is that both parks and homogeneous forests, being more disturbed by human activity (for instance, higher trampling), favour opportunistic or disturbance-tolerant species (Gray 1989), thus boosting species counts.

The composition of carabid beetle communities differed clearly among our three habitat types. Deadwood quantity and canopy cover emerged as two key environmental variables shaping these communities. As expected, parks and forests hosted distinctly different communities due to their contrasting habitats. Carabid beetles are sensitive to habitat type (Lindroth 1985, 1986; Lövei & Sunderland 1996; Kotze et al. 2011), with clear distinctions between open-habitat specialists and forest specialists, a pattern confirmed by our trait analysis showing fewer open-habitat species in forests and fewer forest specialists in parks. An interesting finding was the low variation in community composition within homogeneous forests compared to both heterogeneous forests and parks. Homogeneous forests supported spatially similar and stable communities, whereas heterogeneous forests exhibited much greater variation between sites. One explanation is that, in heterogeneous forests, different microhabitats within the same forest patch may harbour distinct species, even over small spatial scales. Since our traps were clustered in a single location per site, they likely sampled only the species present locally, which may vary across the random sampling point in other heterogeneous forests. In contrast, the uniform microhabitats in homogeneous forests result in similar species compositions regardless of trap location, producing more consistent communities across sites. This spatial consistency further supports the accuracy of our visual assessment of heterogeneity from the beetles’ perspective. Deadwood and canopy openness, variables emphasised in our visual evaluation, were indeed crucial in shaping community composition. Previous studies have similarly demonstrated that carabid communities respond to habitat heterogeneity (Lassau et al. 2005; Thorn et al. 2016) and canopy cover (Thorn et al. 2016), aligning with our findings. Future research focusing on spatial variation within forests is required to provide deeper insights into how microhabitat heterogeneity influences beetle communities at finer scales.

We found fewer macropterous (winged) species in forests compared to parks, and more brachypterous (short-winged) species in homogeneous forests than in heterogeneous ones. This pattern is common in carabid beetles, with winged, more dispersive species typically more abundant in urban parks than in rural or forested environments (Sadler et al. 2006; Niemelä and Kotze 2009), and similarly, more winged species are found in grasslands compared to forests (Ogai and Kenta 2016). Comparable dispersal trends have been observed in other taxa such as spiders (Bonte et al. 2006). Wing polymorphism is likely an adaptive response to habitat fragmentation, facilitating dispersal among suitable habitat patches, whether between isolated parks or within patchy heterogeneous habitats (Hastings 1983; Finand et al. 2024). In contrast, in homogeneous forests where suitable patches are larger and better connected, dispersal ability is less critical. However, this pattern is not universal across all organisms. For example, ants (Finand et al. 2023), butterflies (Schtickzelle et al. 2006), and plants (Cheptou et al. 2008) may show different dispersal strategies when the costs of moving through unsuitable environments are high. Surprisingly, we found no correlation between beetle body size and heterogeneity. Several studies have linked carabid beetle size to environmental factors, for example, higher disturbance often favours smaller species (Blake et al. 1994), urban environments tend to support smaller species than forests (Sadler et al. 2006), and more complex environments are associated with smaller species of darkling beetles (Bartholomew et al. 2016). At the population level, we detected no consistent effect of habitat type on beetle size or mass, except in a few species. A common hypothesis is that individuals developing in warmer environments grow faster as larvae and thus mature at smaller sizes (Brown et al. 2004). A potential explanation for the lack of size differences in our study is that all forests were located within an urban matrix, where temperature differences were relatively small. However, size differences have been reported for species like *Carabus nemoralis* in other studies (Weller and Ganzhorn 2004).

In summary, our study highlights the complex influence of microhabitat heterogeneity on carabid beetle communities within remnant urban forests. While heterogeneity affects community composition, it does not automatically result in an increase in species richness, indicating that factors such as habitat type, disturbance, and species-specific responses also play important roles. Notably, the heterogeneous forests in our study, characterised by higher amounts of deadwood and an open canopy, reflect management practices that retain structural elements critical for biodiversity (see Kotze et al. 2022). This suggests that forest management strategies promoting deadwood retention and canopy openness can enhance habitat quality for many beetle species, even in the urban milieu. The contrasting variability observed between homogeneous and heterogeneous forests further emphasises the need to consider fine-scale habitat structure when assessing biodiversity responses. Future research is needed to explore how different management regimes influence microhabitat features over time and their cascading effects on beetle communities. These studies could investigate genetic adaptation and phenotypic plasticity (see Desender 1989; Sheridan and Bickford 2011) in response to management-induced heterogeneity, thus providing valuable insights into species’ resilience in these highly dynamic urban systems. Overall, integrating ecological knowledge with sustainable forest management is essential to conserve functional and diverse beetle communities in increasingly fragmented landscapes.

## Supporting information

Sup Mat

## Acknowledgements

We thank Valeria Carini for her assistance in the field and laboratory during the summer of 2021, Changyi Lu for his guidance regarding soil analyses, Ida Pohjanlehto for her help with beetle identification and Sampsa Malmberg for identifying some of the more difficult individuals. We are also grateful to Lahti University Campus and the University of Helsinki for offering a start-up grant (MT) for the fieldwork and the City of Lahti for providing access to the study sites. AI was used to improve the English language.

## Statements & Declarations

### Author contributions

Johan Kotze, Heikki Setälä, and Meeri Tahvanainen conceived and designed the study and collected the data. Basile Finand performed the analyses. The first draft of the manuscript was written by Basile Finand. All authors commented on previous versions of the manuscript. All authors read and approved the final manuscript.

### Data availability

The data and codes will be available on Zenodo if the paper is accepted.

