## Supplementary material for "Perceived urban microhabitat heterogeneity impacts carabid beetle communities": Sup Mat

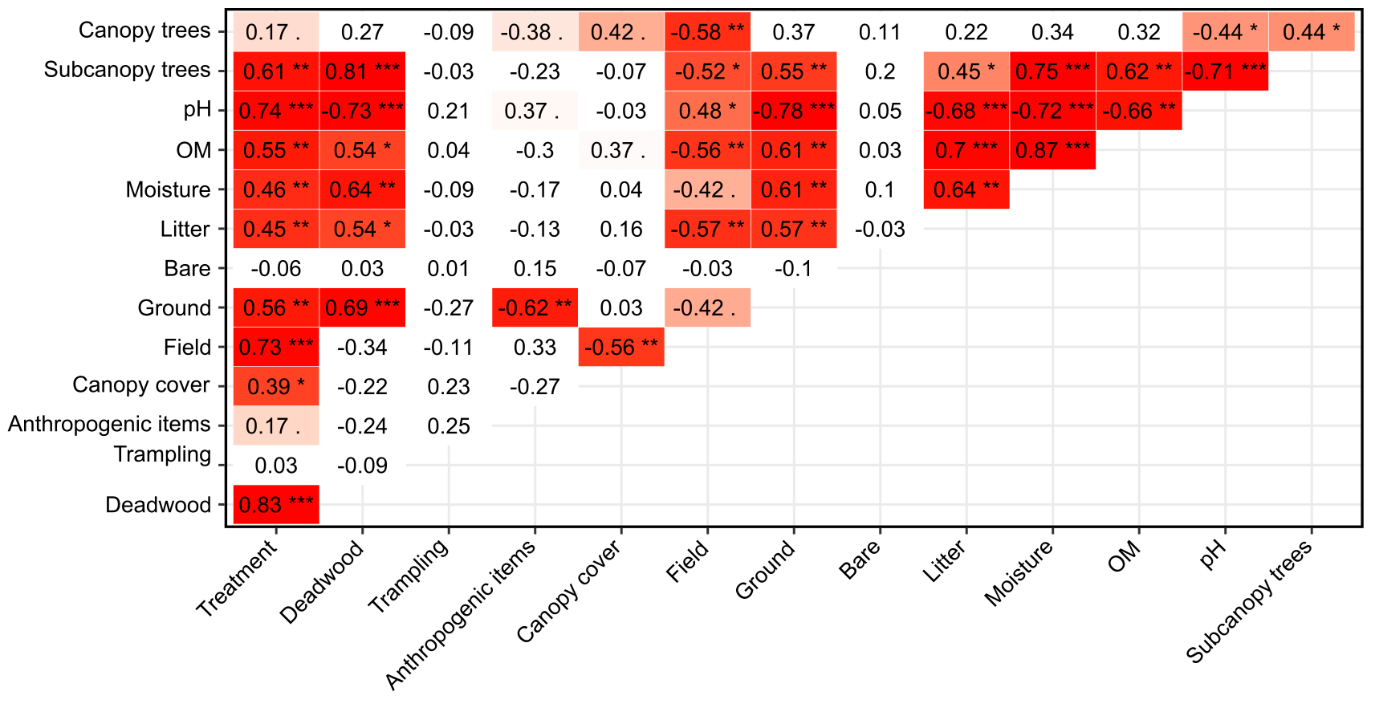

**Fig. S1**: Correlation heatmap between all the environmental variables measured in this study. Values in each box are the effect size (rho for the Spearman tests, and epsilon2 for the Kruskal-Wallis tests). Dark red indicates a p-value < 0.05, and light red a p-value < 0.1. Three stars indicate a p-value < 0.001, two stars a p-value < 0.01, one star a p-value < 0.05, and a dot a p-value < 0.1.

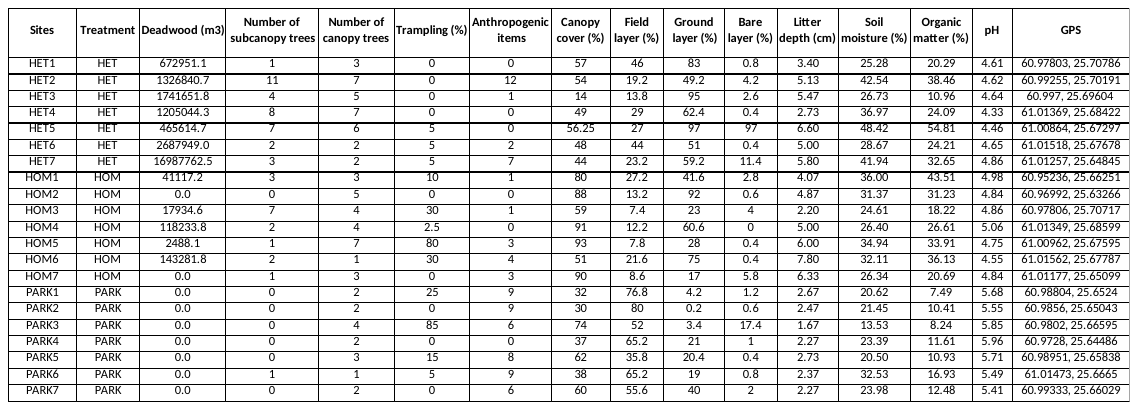

**Table S1**: Environmental characteristics measured for each site and coordinates. HET = heterogeneous forests, HOM = Homogeneous forests, and PARK = urban parks. Canopy cover, Field layer, Ground layer, Bare Layer, and Litter depth are means (see Methods).

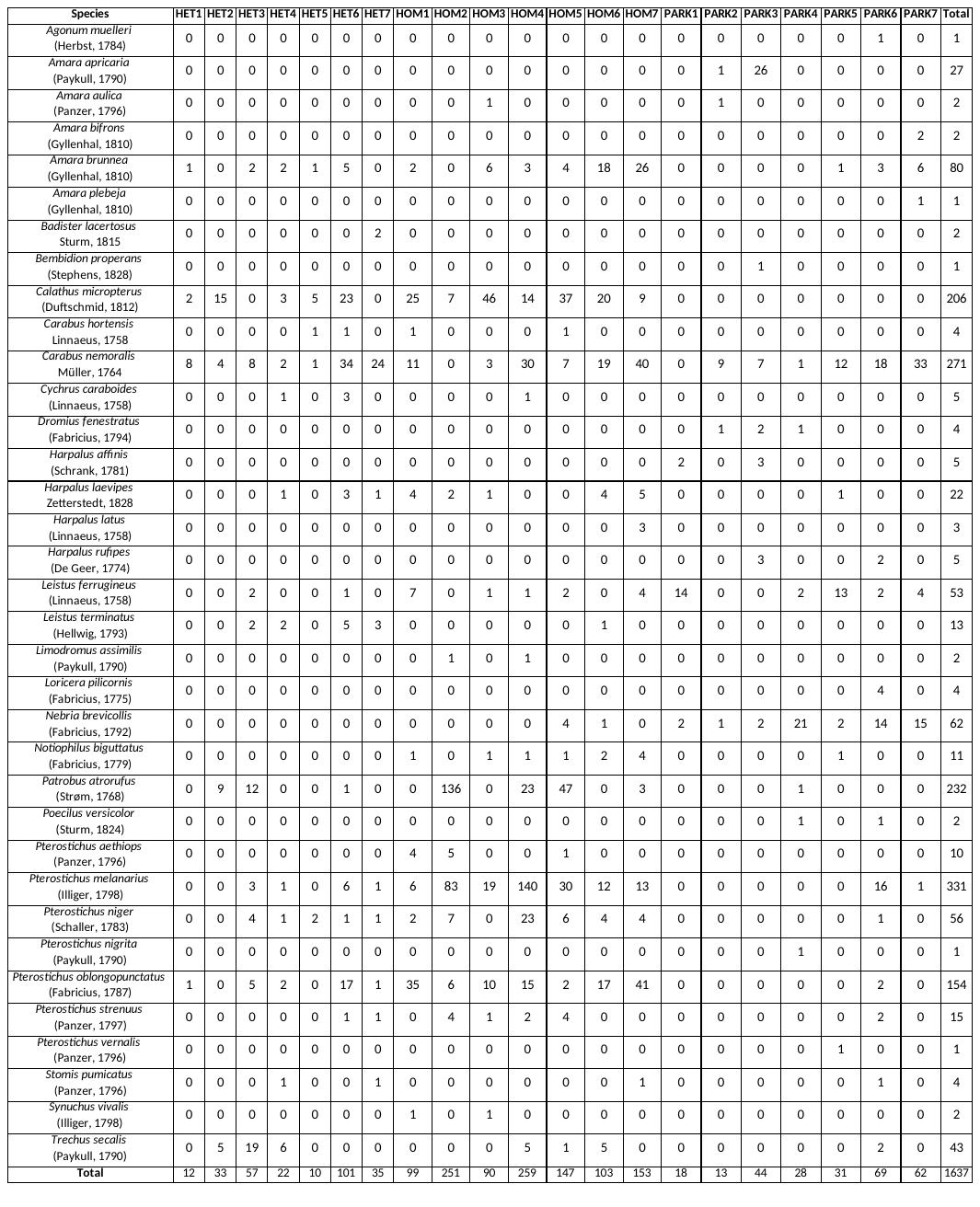
**Table S2:** List of species per site. The values represent the number of individuals per species collected from this study.
